# Temporal Development of the Tracheal Microbiome Across Production Phases in Broiler Chickens

**DOI:** 10.64898/2026.07.11.737896

**Authors:** Seif Hundam, Mohammad Borhan Al-Zghoul, Raghad Alomari, Sandra Nammas, Mu’taz Almaasfeh, Hebah Aboomer, Shirin Qaaty, Shahed Ogiliat, Duha Makableh, Shadi Shahatit, Ghaith Alhamouri

## Abstract

The respiratory microbiome plays important roles in poultry health, immune regulation, and pathogen resistance, yet its development throughout the broiler production cycle remains poorly understood. This study investigated temporal changes in the tracheal microbiome of broiler chickens across production phases. Tracheal samples were collected during the starter (day 12), grower (day 21), early finisher (day 26), and late finisher (day 35) phases and analyzed using 16S ribosomal RNA gene sequencing. Tracheal microbial richness, diversity, community structure, and taxonomic composition changed significantly across broiler production stages, including starter, grower, early finisher, and late finisher feeding phases. Alpha diversity increased progressively throughout production, with significant increases in richness, diversity, and phylogenetic diversity during later stages. Beta diversity analysis revealed distinct microbial communities associated with each production phase, with starter-phase samples clearly separated from later phases. Taxonomic profiling showed dominance of Proteobacteria during the starter and grower phases, with enrichment of *Methylobacterium-Methylorubrum* and *Pseudomonas* during the starter phase and of *Escherichia-Shigella* during the grower phase. In contrast, the finisher phases exhibited reduced Proteobacteria abundance and increased Firmicutes and Actinobacteriota, including *Lactobacillus*, *Ligilactobacillus*, *Faecalibacterium*, *Streptococcus*, *Staphylococcus*, *Romboutsia*, and *Corynebacterium*. Overall, the tracheal microbiome underwent progressive maturation, shifting from a Proteobacteria-dominated community to a more diverse, complex, Firmicutes-rich ecosystem. These findings provide new insights into the development of the respiratory microbiome in broiler chickens and may support strategies to improve poultry respiratory health. Because dietary transitions occurred concurrently with age progression, the observed microbiome shifts should be interpreted as production-stage-associated changes rather than diet-specific effects.

## Introduction

The term “microbiota” refers to the microbial community, which includes commensal, symbiotic, and pathogenic microorganisms that typically inhabit a region of an animal and are approximately twice as abundant as the host’s somatic and germinal cells <u>(1)</u>. In broiler chickens, the gut microbiome has been extensively studied due to its critical roles in immune regulation, nutrient metabolism, and gut physiology. Additionally, the microbiota contributes to host defense by limiting colonization of enteric pathogens through competitive exclusion and the production of bacteriostatic and bactericidal compounds <u>(2)</u>. While much of this research has focused on the gut, emerging evidence suggests that similar microbial-mediated mechanisms operate in other mucosal systems, including the respiratory tract.

Respiratory diseases remain one of the leading causes of economic losses in the poultry industry <u>(3–5)</u>. Increasing evidence indicates that the respiratory tract microbiome plays a pivotal role in maintaining respiratory health in birds <u>(6,4)</u>. This microbial community contributes to immunocompetence and protects against pathogen colonization through mechanisms similar to those observed in the gut <u>(7–9)</u>. The diversity and composition of the respiratory microbiota are influenced by multiple factors, including intrinsic host characteristics (e.g., breed, age, and genetics) and extrinsic factors (e.g., production systems, diet, antibiotic use, environmental conditions, and litter quality) <u>(10)</u>.

The trachea, a key component of the avian respiratory system, functions not only as a conduit but also as an important site of immune surveillance. Its epithelial lining, mucociliary apparatus, and associated lymphoid tissues provide a first line of defense against inhaled pathogens <u>(2,9)</u>. The tracheal microbiota—a diverse assemblage of bacteria, fungi, and viruses—resides in this environment and helps maintain respiratory homeostasis. Advances in high-throughput sequencing have highlighted the importance of this microbial ecosystem, demonstrating that it can increase susceptibility to respiratory diseases <u>(11)</u>.

Diet is one of the most influential factors shaping microbial communities in animals. In poultry production systems, birds undergo distinct production phases—starter, grower, and finisher—each formulated to meet changing nutritional requirements during development. For example, starter diets with higher nutrient density have been associated with increased populations of beneficial bacteria, such as *Lactobacillus* and *Bifidobacterium,* in the gut. At the same time, dietary changes can differentially affect microbial communities across intestinal regions <u>(13)</u>. Furthermore, diet has been recognized as a key driver of microbial complexity and community structure in the gut <u>(14)</u>.

Despite these insights, the extent to which production phase transitions influence the tracheal microbiome remains poorly understood. Although previous studies have described respiratory microbiota composition in broilers across age, health status, environmental stress, or production systems, few have examined tracheal microbiome succession at defined commercial feeding-phase transitions using adequately replicated sampling. Such dietary effects may alter microbial diversity, community structure, and the relative abundance of key taxa, with potential implications for host health and productivity. Therefore, the present study addresses this specific gap by characterizing the development of the tracheal bacterial community across key production stages corresponding to starter, grower, early finisher, and late finisher diets. By examining microbial shifts across multiple developmental time points, this study seeks to provide novel insights into the temporal dynamics and potential production-stage-associated modulation of the avian respiratory microbiome, ultimately supporting strategies to improve respiratory health and optimize poultry performance.

## Materials and methods

The Animal Care and Use Committee of Jordan University of Science and Technology (JUST-ACUC) reviewed and approved all procedures conducted in the current study (Approval Number:16/4/12/6).

### Study population and incubation

In total, 1000 fertile Ross 308 broiler eggs were obtained from a single commercial breeder flock to minimize genetic variability. These eggs were weighed, and only those within the 50–56 g weight range were selected for use in the experiments. Eggs with obvious defects or abnormalities, such as cracks or irregular shapes, were excluded. The selected eggs were randomly allocated to four commercial Type I HS-SF incubators (Masalles Company, Inc., Barcelona, Spain). All eggs were automatically turned every hour throughout incubation <u>(15)</u>.

On ED 7, all eggs were candled to identify and remove infertile eggs and those containing non-viable embryos, thereby minimizing the risk of contamination from decomposing eggs during incubation. Eggs were incubated under standard commercial conditions at 37.5°C and 56% relative humidity (RH) during the first 19 days of incubation. From embryonic day (ED) 19 to ED 21, the temperature was reduced to 37.0°C, the relative humidity increased to 65%, and egg turning was discontinued in all incubators.

### Hatching management and post-hatching rearing

On the day of hatching, chicks were kept in the incubator until they were completely dry. One-day-old chicks were then transferred to the Animal House at the Jordan University of Science and Technology to initiate the field study. Chicks were randomly allocated to cage pens (8 chicks per cage pen), with each cage pen considered an experimental unit.

The chicks were reared for 35 days under standard husbandry conditions <u>(16)</u>. The ambient temperature was initially maintained at 35 °C during the first week, with continuous lighting (23 h light: 1 h dark), and was gradually reduced to 21 °C by day 24. Feed and water were provided ad libitum throughout the study.

The chicks were fed corn–soybean–wheat-based diets formulated according to National Research Council recommendations, consisting of three feeding phases (**Table 1**): Starter (days 1–12), Grower (days 13–24), and Finisher (days 25–35) <u>(17)</u>. These diets provided metabolizable energy (ME) levels of 2896, 3008, and 3175 kcal/kg, respectively, and crude protein (CP) levels of 21.6%, 19.4%, and 18.1%, respectively.

**Table 1.** Ingredient composition of experimental starter, grower, and finisher diets (% of diet).

| Ingredient (% of Diet) | Starter | Grower | Finisher |
| --- | --- | --- | --- |
| Corn | 47.90 | 51.30 | 50.20 |
| Soybean meal | 37.00 | 31.00 | 28.80 |
| Wheat | 10.00 | 12.50 | 15.00 |
| Oil | 0.80 | 1.30 | 2.50 |
| Calcium Carbonate | 1.30 | 1.20 | 1.15 |
| MCP | 0.85 | 0.60 | 0.38 |
| Salt | 0.26 | 0.28 | 0.25 |
| Sodium Bicarbonate | 0.15 | 0.14 | 0.13 |
| Lysine sulphate | 0.63 | 0.55 | 0.53 |
| Methionine | 0.37 | 0.35 | 0.35 |
| Threonine | 0.16 | 0.14 | 0.14 |
| Valine | 0.06 | 0.05 | 0.06 |
| Arginine | 0.01 | 0.02 | 0.02 |

| **Ingredient (% of Diet)** | **Starter** | **Grower** | **Finisher** |
| --- | --- | --- | --- |
| Premix Vitamin 0.1% | 0.10 | 0.10 | 0.10 |
| Min. Premix 0.1% | 0.10 | 0.10 | 0.10 |
| Choline Chloride | 0.08 | 0.08 | 0.08 |
| Phytase Enzyme | 0.02 | 0.02 | 0.02 |
| NSP Enzyme | 0.01 | 0.01 | 0.01 |
| Essential Oils | 0.01 | 0.01 | 0.01 |
| Toxin Binder | 0.16 | 0.17 | 0.17 |
| Butyric acid | 0.05 | 0.05 | 0.05 |
| **Total** | **100.00** | **100.00** | **100.00** |

### Microbiological analysis

#### Sample collection

Samples were collected on post-hatch days (D) 12, 21, 26, and 35, representing key transitions between production phases: the starter phase (1S; D12), grower phase (2G; D21), early finisher phase (3FE; D26), and late finisher phase (4FL; D35). At each time point, one chick was randomly selected from each of the 24 replicate pens per experimental group (n = 24 per group). Each pen was considered an experimental unit for statistical analysis. Selected birds were fasted for 12 h before sampling to minimize crop content contamination, then humanely euthanized by cervical dislocation and sampled.

The trachea was aseptically removed from each selected chick using sterile instruments and immediately snap-frozen on-site with liquid nitrogen to prevent DNA degradation. Samples were stored in a CryoCube F570 Series Ultra-Low Temperature (ULT) Freezer (Eppendorf, Hamburg, Germany) at −80°C to preserve the integrity of the microbial DNA until extraction.

#### DNA Isolation and Sequencing

The tracheal lumen was lavaged with the CD1 solution provided in the DNeasy PowerSoil Pro kit, and subsequent isolation steps were performed according to the manufacturer’s instructions (QIAGEN, June 2023). The hypervariable V3-V4 region of the 16S rRNA gene was amplified with the universal primers Bakt_341F: (CCTACGGGNGGCWGCAG) and Bakt_805R: (GACTACHVGGGTATCTAAT CC). Sequencing was performed commercially by Microsynth AG, Balgach, Switzerland, using an Illumina MiSeq platform (Illumina, CA, USA) following the 2 × 300 bp paired-end sequencing protocol.

#### Data Analysis by Bioinformatic Tools

Bioinformatic processing was conducted using QIIME 2 (version 2025.7) <u>(18)</u>. Sequence quality control, noise reduction, and the construction of amplicon sequence variants (ASVs) were performed using the DADA2 plugin. Taxonomic assignment was based on the SILVA 138 reference set, which contained 99% of the operational taxonomic units (OTUs). To ensure comparability, all datasets were rarefied to 41500 reads per sample before calculating diversity indices. Measures of alpha diversity (ace, shannon, and Faith’s Phylogenetic Diversity) were determined using the Kruskal–Wallis test implemented in QIIME 2. For community structure ordination, Principal Coordinates Analysis (PCOA) was performed using the MicrobiomeStat package in R. Differences in microbial community composition (beta diversity) among groups were evaluated using PERMANOVA with Bray–Curtis distances. Homogeneity of multivariate dispersions was validated using PERMDISP, which revealed no significant differences in community variance across the four groups (F = 0.23, p = 0.84, 999 permutations), confirming that any observed compositional differences were due to centroid shifts rather than unequal group dispersions. Differential abundance analysis (DAA) was conducted using the linear models for differential abundance analysis (LinDA) method from the MicrobiomeStat R package. Taxa with an FDR-adjusted P value less than 0.05 were considered differentially abundant. To prevent potential laboratory or reagent-kit contamination from biasing biological interpretations, statistical decontamination was performed on the raw, unrarefied sequence table using the decontam frequency model in QIIME 2. Out of 2,389 initial features, 7 were identified as statistical contaminants and removed, while 2,382 non-contaminant features were retained for downstream analysis. The comprehensive quality filtering profile, including individual feature P-scores, raw abundances, and prevalence metrics, has been provided as an interactive QIIME2 supplementary artifact (Supplementary Material File S4).

## Results

### Effects of the production phases on the microbial alpha diversity

To evaluate the temporal impact of production phases on the microbial richness, evenness, and phylogenetic diversity within the trachea, alpha diversity indices—specifically the ACE index, Shannon index, and Faith’s Phylogenetic Diversity index—were analyzed across the four production periods: starter (1S), grower (2G), early finisher (3FE), and late finisher (4FL). Significant variations in microbial structure were observed as the production phases progressed, with a general trend toward increased diversity and richness over time **Figure 1**.

**Figure 1.**
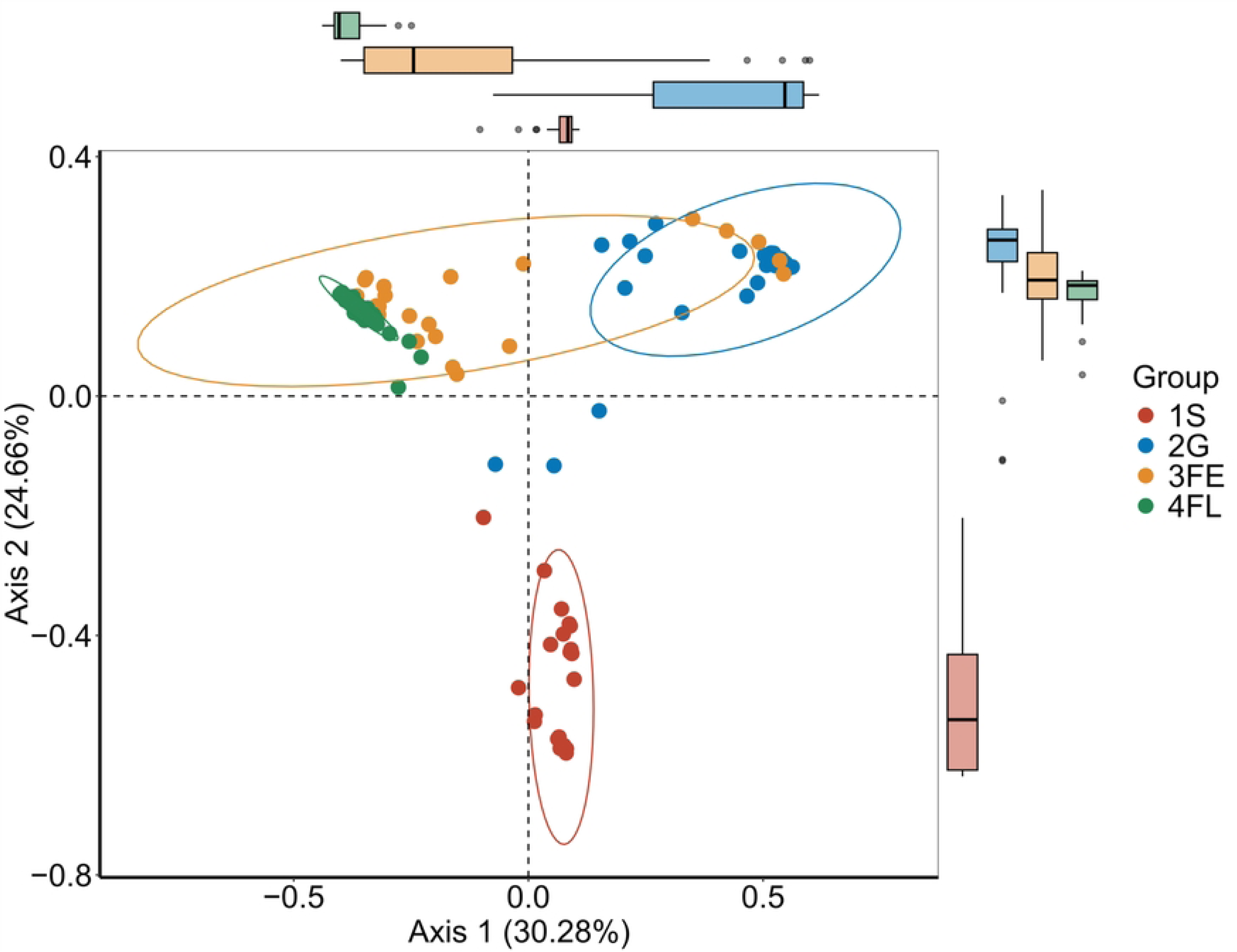
Alpha diversity indices of the tracheal microbiota are depicted in box plots, illustrating the impact of production phases on tracheal microbial diversity. Plots display community (a) richness (ACE index), (b) diversity and evenness (Shannon index), and (c) phylogenetic diversity (Faith’s PD index). Different letters above the bars signify statistically significant differences (q < 0.05) among the groups. The groups are categorized as follows: Starter (1S), Grower (2G), Early Finisher (3FE), and Late Finisher (4FL). The sample size (n) was 24 per group.

Microbial community richness, as assessed by the ACE index, showed a clear, step-by-step increase across all production phases, rising from a baseline during the starter phase (1S) to the highest community richness in the late finisher phase (4FL) (all pairwise comparisons, raw p < 0.05, FDR-adjusted q < 0.05, Supplementary Table 1). This trend was closely mirrored by overall community diversity and evenness (Shannon index), which expanded from 1S through 2G, culminating in a highly elevated, tightly clustered diversity plateau by the 4FL phase (all pairwise comparisons, raw p < 0.05, FDR-adjusted q < 0.05). Concurrently, the phylogenetic diversity of the microbiota (Faith’s PD index) remained statistically static during the initial transition from the 1S to the 2G phase (FDR-adjusted q > 0.05), followed by a significant, non-linear expansion during the early finisher (3FE) phase (raw p < 0.05, FDR-adjusted q < 0.05) and achieving peak phylogenetic diversity in the 4FL cohort (raw p < 0.05, FDR-adjusted q < 0.05).

### Effects of the production phases on the microbial beta diversity

To evaluate the impact of production phases on the overall structural similarity of beta diversity in the tracheal microbiome, PCOA was performed using a Bray-Curtis distance matrix. Ordination analysis revealed a distinct, phase-dependent spatial segregation of the microbial communities across the four production phases (**Figure 2**). PERMANOVA confirmed that microbial community composition differed significantly among production phases (q < 0.001). The starter phase (1S) formed a highly segregated, independent cluster along the negative region of Axis 2, establishing a distinct baseline community. Transition to the grower diet (2G) induced a major shift toward the upper right quadrant. As animals advanced to the early finisher phase (3FE), the community exhibited heightened dispersion, partially overlapping with adjacent phases. However, by the late finisher phase (4FL), the tracheal community stabilized into an exceptionally tight, highly uniform cluster on the far left of Axis 1 (all pairwise PERMANOVA comparisons, raw p < 0.001, FDR-adjusted q < 0.001, Supplementary Table 2).

**Figure 2.**
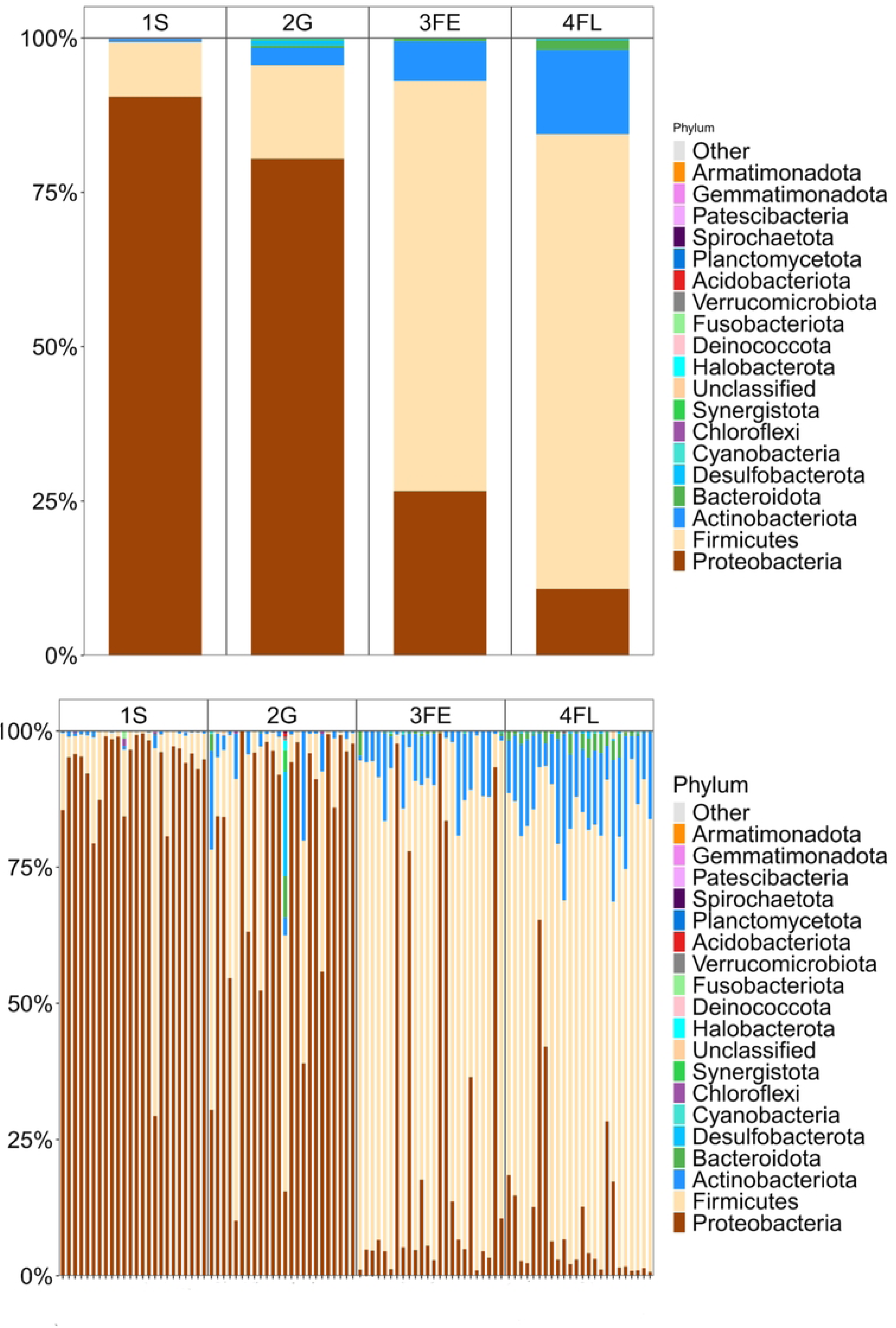
Principal Coordinates Analysis (PCOA) of beta diversity based on the Bray-Curtis dissimilarity is presented. ordination plot mapping the beta diversity shifts across the Starter (1S), Grower (2G), Early Finisher (3FE), and Late Finisher (4FL) production phases. The box plots at the top and right of the main panel depict the distribution of samples along Axis 1 and Axis 2, respectively. Significant differences in microbial community composition were detected among production phases (PERMANOVA q < 0.001). In all panels, ellipses represent the 95% confidence interval for each group. The percentage of variation explained by each axis is presented in parentheses.

### Differential abundance analysis

Taxonomic profiling revealed a highly dynamic, phase-specific restructuring of the tracheal microbiome across the feeding timeline, characterized by a distinct transition from early Proteobacteria dominance toward a Firmicutes-rich, highly even community (**Figures 3** and **4**).

**Figure 3.**
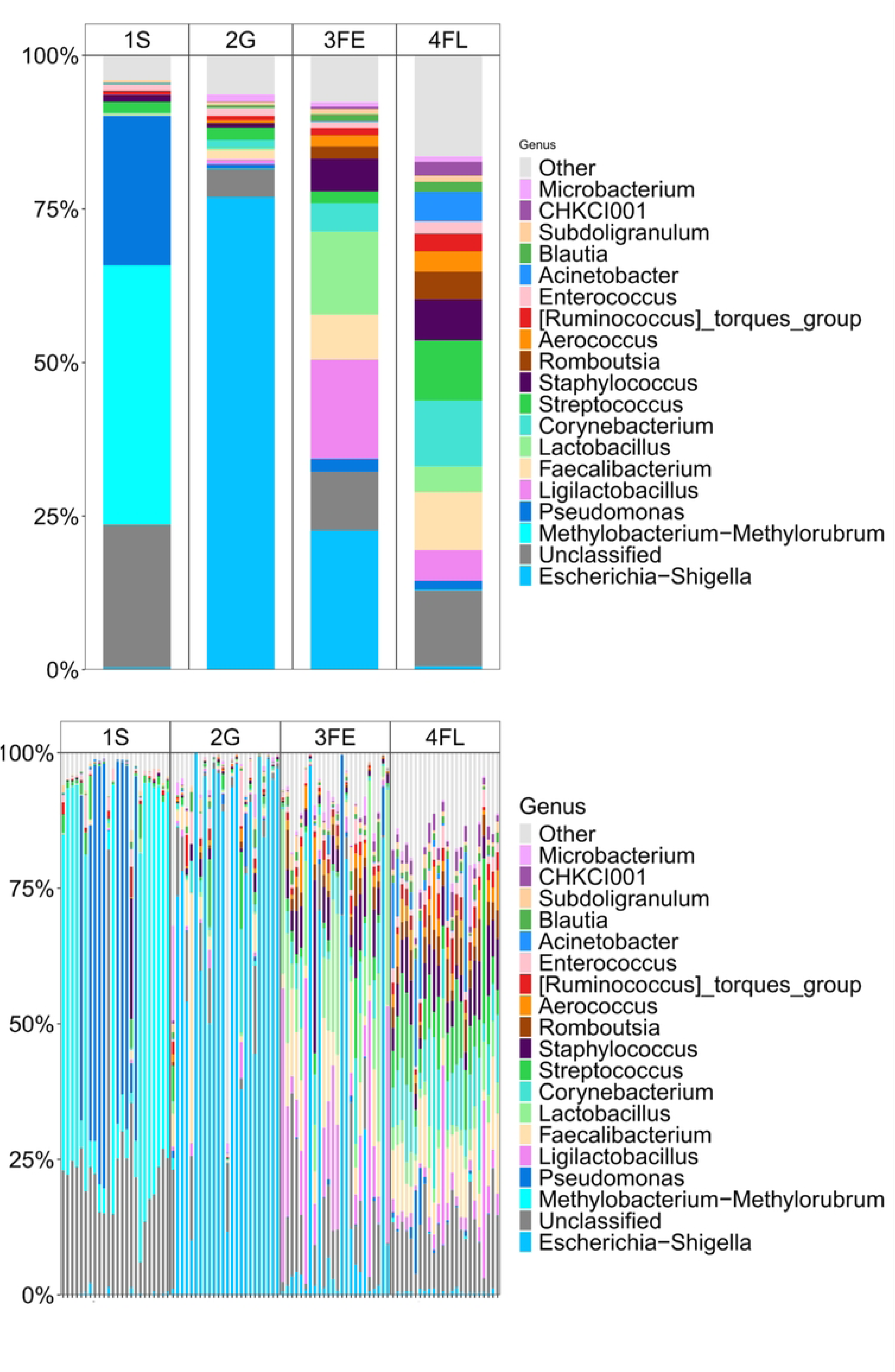
Relative taxonomic abundance profiles of major bacterial phyla across the four production phases (1S, 2G, 3FE, and 4FL), highlighting the compositional inversion from Proteobacteria to Firmicutes. (Top) stacked bar plot showing the mean relative abundance of the major bacterial phyla across all groups. (Bottom) stacked bar plot showing the relative abundance of the major bacterial phyla for each sample.

**Figure 4.**
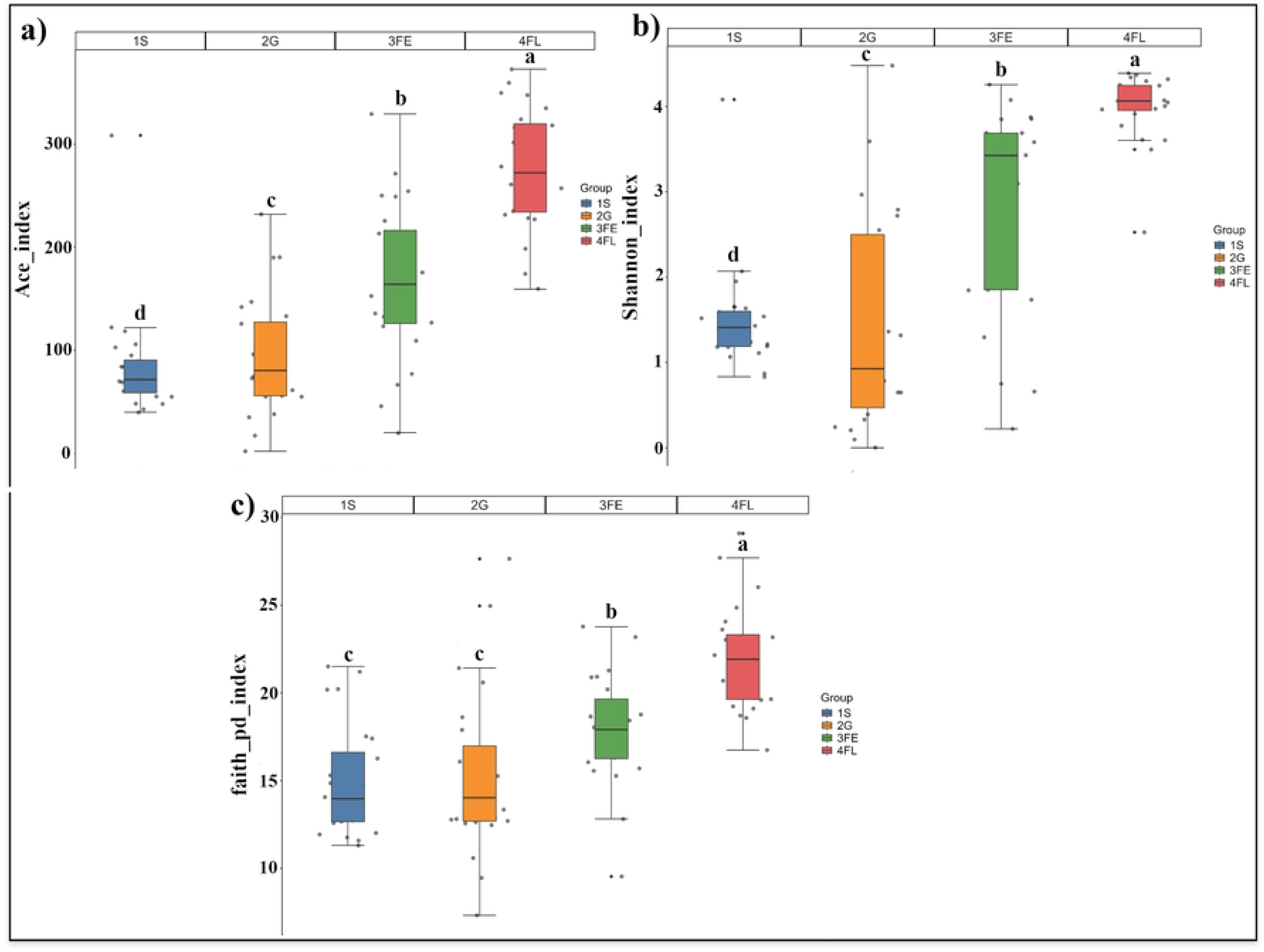
Relative abundance of microbial genera across the four production phases (1S, 2G, 3FE, and 4FL), highlighting the transition from early-stage genera (Methylobacterium-Methylorubrum, Acinetobacter, and Escherichia-Shigella) to late-stage genera (Corynebacterium, Streptococcus, Staphylococcus, and Romboutsia). **(Top) stacked** bar plot showing the mean relative abundance of the major bacterial genera across all groups. (Bottom) stacked bar plot displaying the relative abundance of the main bacterial genera for each sample.

At the phylum level, the starter (1S) and grower (2G) phases were significantly enriched with Proteobacteria (>80%; LinDA q < 0.01; Supplementary Table 3). This pattern was mainly due to an initial abundance of *Methylobacterium-Methylorubrum* and *Pseudomonas* in the 1S phase, which shifted to enrichment of *Escherichia-Shigella* during the 2G period (LinDA q < 0.01; **Figures 3** and **4**).

The early finisher (3FE) phase was linked to a decrease in Proteobacteria, along with an increase in Firmicutes and Actinobacteriota (LinDA q < 0.001; Figure 3), which occurred alongside the enrichment of specific anaerobic commensals and environmentally derived genera, such as *Ligilactobacillus*, *Faecalibacterium*, and *Lactobacillus* (LinDA q < 0.01); **Figure 4**). During the late finisher (4FL) phase, community evenness reached its peak, and Proteobacteria were found at a minimal level, making room for a balanced, diverse group of taxa—including *Corynebacterium*, *Streptococcus*, *Staphylococcus*, and *Romboutsia*—along with an increase in lower-abundance taxa grouped under “Other” (**Figures 3** and **4**). Overall, these compositional changes indicate a progressive transition from an early Proteobacteria-enriched community to a more diverse, taxonomically complex respiratory microbiome during later production stages.

## DISCUSSION

The present study investigated the temporal dynamics of the tracheal microbiome of broiler chickens across different production phases. Our results show that these production phases are associated with significant changes in microbial richness, diversity, community structure, and taxonomic composition. Overall, the findings align with observed temporal shifts in the tracheal microbiome during commercial production, marked by increasing microbial complexity and a clear transition from an early Proteobacteria-dominated community to a more diverse and richer microbial ecosystem.

A major finding of this study was the progressive increase in alpha diversity indices across the feeding phases. Both the ACE and Shannon indices increased significantly from the starter to the late finisher phase, indicating a gradual expansion in microbial richness and community evenness. Similarly, Faith’s phylogenetic diversity increased during later developmental stages, suggesting the recruitment of phylogenetically distinct taxa over time. These findings are consistent with observations from ileal, caecal, and lung microbiome studies in broiler chickens, where microbial communities become increasingly diverse as birds mature and are exposed to a wider range of environmental and dietary stimuli <u>(19,20)</u>.

The observed increase in tracheal microbial diversity may reflect the combined influence of age-related host development and dietary transitions. As birds progress through production stages, changes in nutrient composition, feed particle size, feeding behavior, and environmental exposure may alter the tracheal environment, thereby facilitating colonization by additional microbial taxa. While previous studies have associated greater microbial diversity with enhanced community resilience and competitive exclusion of pathogens Larsen et al., (<u>21);</u> Frances et al., <u>22)</u>, Abuin-Denis et al. <u>(23)</u> recently challenged this paradigm, reporting that colonization resistance was better explained by microbial network integrity than by diversity alone. Because the present study did not evaluate microbial interaction networks, the functional implications of the observed increases in diversity and evenness remain uncertain. Therefore, although the mature tracheal microbiome exhibited greater diversity and a more even taxonomic structure, we cannot determine whether these characteristics confer enhanced colonization resistance or ecosystem stability. Future studies incorporating co-occurrence network analyses, microbial interaction modeling, and functional metagenomic approaches will be necessary to determine whether tracheal microbiome maturation involves not only increased diversity but also the establishment of robust microbial interaction networks that contribute to respiratory health and pathogen resistance.

Microbial community structure differed markedly among production phases. The distinct clustering observed in PCOA ordination, together with highly significant PERMANOVA results, indicates that each feeding phase was characterized by a unique microbial assemblage. The clear separation of the starter-phase samples from subsequent phases suggests that the early-life tracheal microbiome represents a relatively immature community that undergoes substantial restructuring during post-hatch development. The greater dispersion observed during the early finisher phase may indicate a transitional stage during which microbial succession is actively occurring. In contrast, the tight clustering observed during the late finisher phase suggests greater similarity and stability among individuals.

This pattern of phase-specific clustering is consistent with the findings of Shen et al. <u>(24)</u>, who reported that “obvious and regular changes were observed in the Beta diversity of the pulmonary microbiota in broilers during the growth cycle.” Our results indicate that respiratory microbial communities undergo substantial compositional restructuring as birds mature. However, unlike the present study, Shen et al. observed no significant changes in alpha diversity throughout the growth cycle, whereas we detected progressive increases in the diversity. Several factors may explain this discrepancy. First, the studies examined different respiratory niches: Shen et al. focused on the lung microbiota, whereas the present study investigated the tracheal microbiota. Distinct anatomical locations within the respiratory tract harbor unique microbial communities due to differences in local environmental conditions, immune activity, and microbial exposure. Second, our study specifically evaluated microbiome dynamics across defined production phases, whereas Shen et al. did not evaluate microbiome dynamics across defined production-phase transitions. Third, differences in sample size may have influenced the statistical power to detect diversity changes. Shen et al. analyzed six lung samples per sampling point. In contrast, the present study included 24 tracheal samples per time point, providing greater power to detect subtle but biologically relevant differences in microbial diversity.

Taxonomic analyses revealed pronounced compositional shifts throughout the study period. During the starter and grower phases, the tracheal microbiome was dominated by Proteobacteria, followed by a marked shift toward a Firmicutes-rich community during the finisher phases. This pattern aligns with a meta-analysis by Moreno-León et al. <u>(25)</u>, which reported that tracheal colonization by Firmicutes was accompanied by a decline in Proteobacteria and a progressive increase in Firmicutes abundance. Moreno-León et al. explained this early Proteobacteria dominance by noting that these taxa are already established in the respiratory microbiota before hatching, consistent with our observation of the highest Proteobacteria abundance in starter-phase samples. The subsequent increase in Firmicutes observed during later production stages may reflect the combined effects of dietary transitions, host developmental maturation, and cumulative environmental exposure, all of which are known to influence microbial community assembly in poultry.

The enrichment of genera such as *Methylobacterium-Methylorubrum* and *Pseudomonas* during the starter phase may reflect environmental acquisition from hatchery and housing environments. *Methylobacterium-Methylorubrum* was detected in bronchoalveolar lavage and lung tissue, suggesting that it is a part of the respiratory microbiota <u>(26,27)</u>. Notably, *Pseudomonas* in the respiratory microbiota was detected exclusively during the starter phase in our study, suggesting rapid environmental acquisition followed by displacement. A similar pattern of early *Pseudomonas* predominance was observed by Mulholland et al. <u>(11)</u>, who reported that this genus was not found at significant levels after Week 1. This temporal pattern characterizes *Pseudomonas* as an easily replaceable taxon, as it was detected only during the starter phase at significant levels, with a substantial decrease in later phases.

The expansion of *Escherichia-Shigella* during the grower phase may indicate continued microbial recruitment from feed, litter, water, or bird-to-bird interactions, as *Escherichia-Shigella* is among the core genera in the chicken respiratory microbiome <u>(28)</u>. Our findings align with evidence that high *Escherichia-Shigella* loads in the trachea represent transient colonization rather than permanent community establishment. According to <u>(29)</u>, despite high *Escherichia-Shigella* dominance at week 3, the tracheal microbiota shifted to *Gallibacterium* by the end of the fattening cycle, which they interpret as evidence that mucociliary and immune clearance mechanisms alter the tracheal epithelial environment, reducing *Escherichia-Shigella* colonization. The transient nature of *Escherichia-Shigella* enrichment supports our observation that the decline in early Proteobacteria-enriched taxa during later production stages may be consistent with age-related changes in the tracheal environment, including immune and mucociliary maturation; however, these mechanisms were not directly measured in the present study.

Several Firmicutes genera enriched during the later production stages, including *Lactobacillus*, *Ligilactobacillus*, *Faecalibacterium*, and *Romboutsia*, have previously been associated with host health and immune regulation in poultry and other animal species. Although their functional roles within the avian respiratory tract remain incompletely understood, these taxa have been linked to beneficial host–microbiota interactions in various biological systems. For example, *Lactobacillus* and *Ligilactobacillus* have been associated with immunomodulatory activity and protection against respiratory trac tmicrobial infections in poultry and other animals <u>(30–32)</u>. In addition, *Faecalibacterium* has been detected in the broiler trachea and has been reported to correlate with IL-1β expression, suggesting a potential association with local inflammatory responses <u>(9)</u>. Similarly*, Romboutsia* has been positively associated with cecal short-chain fatty acid concentrations and negatively associated with colonic IL-1β levels in mice <u>(33)</u>. While these findings suggest that Firmicutes-enriched communities may possess functional attributes relevant to host health, the present study was limited to taxonomic characterization. It cannot determine whether these genera exert comparable effects within the chicken respiratory tract.

The increase in *Corynebacterium*, *Streptococcus*, and *Staphylococcus* during the late finisher phase is particularly noteworthy, as these genera are commonly detected in the healthy respiratory microbiota of poultry. Johnson et al. <u>(28)</u> identified Corynebacterium and Staphylococcus as core members of the respiratory microbiota. They reported positive associations between these genera and body weight in broilers, whereas Streptococcus showed a negative association. Similarly, Sohail et al. <u>(7)</u> reported higher abundances of Corynebacteriaceae, Staphylococcaceae, and Streptococcaceae in control birds compared with heat-stressed birds, whose respiratory microbiota were dominated by Lactobacillaceae. Collectively, these findings are consistent with previous studies suggesting that variation in respiratory microbiota composition may be associated with broiler health and performance. The concurrent expansion of these taxa in the late-finisher phase, together with numerous low-abundance genera represented within the “Other” category, indicates that the late-finisher tracheal microbiome exhibits greater taxonomic diversity and a more complex community structure than earlier production phases.

The findings of this study have important implications for poultry health and management. Increasing evidence suggests that respiratory microbiota contribute to host defense, immune regulation, and resistance against respiratory pathogens. Therefore, understanding how microbial communities develop throughout production may facilitate the design of nutritional and management strategies to promote beneficial respiratory microbiota. Such approaches contribute to improved respiratory health, enhanced production performance, and reduced reliance on antimicrobial interventions.

However, several considerations should be noted when interpreting this study’s findings. First, dietary phase transitions occurred simultaneously with advancing age—the natural production protocol for broiler chickens. While this prevents isolation of diet-specific effects from age-related development, it enhances the ecological validity of our findings by capturing real-world commercial production conditions. Our observed microbial shifts therefore reflect the integrated influence of dietary changes, age-related physiological development, immune maturation, and cumulative environmental exposure—precisely the combination of factors that drive tracheal microbiome maturation in commercial poultry. This design strength ensures that our findings are directly applicable to poultry management practices, where interventions must address these combined influences rather than isolated factors. Second, this study used repeated cross-sectional sampling, in which different birds were sampled at each time point rather than longitudinal monitoring of the same individuals. Consequently, some variation among sampling periods may reflect inter-individual biological differences. Finally, 16S rRNA gene sequencing provides valuable taxonomic information but does not directly assess the functional activity or metabolic capabilities of the microbial community. Future studies incorporating controlled dietary interventions independent of age, longitudinal sampling designs, and shotgun metagenomic or metatranscriptomic approaches would help clarify the specific mechanisms underlying tracheal microbiome development and function in broilers.

## Conclusion

In summary, our findings suggest that the tracheal microbiome of broiler chickens exhibits marked temporal changes across production phases, characterized by increased diversity, significant restructuring of community composition, and a progressive shift from Proteobacteria dominance toward a more complex, Firmicutes-rich community. These findings provide novel insights into the maturation of the avian respiratory microbiome and establish a foundation for future investigations examining the interactions among diet, host development, respiratory microbial ecology, and poultry health.

## Acknowledgments

The authors extend their sincere gratitude to the Deanship of Research at Jordan University of Science and Technology for providing financial support for this research project. Grant numbers 85/2026 and 68/202 were instrumental in facilitating this work. Additionally, they express their sincere appreciation to Amany Al-Rashadieh and Eng. Ibrahim Alsukhni for their invaluable insights and exceptional technical assistance.

## Data availability

QIIME 2 artifacts, including quality control files, feature tables, taxonomy tables, and rooted phylogenetic trees, associated with this study are publicly available at https://github.com/saifhandam/Tracheal_microbiome_diet.git (accessed on June 17, 2026). Additionally, the raw sequence data are available in the NCBI SRA database under accession PRJNA1479056.

## Supporting information

**S1 Table. Alpha diversity statistics.** This table provides the results of the ACE index comparisons across production phases, including pairwise comparisons, raw p-values, and FDR-adjusted q-values.

**S2 Table. Beta diversity analysis.** This table presents the results of the PERMANOVA analysis for all pairwise comparisons among production phases, including raw p-values and FDR-adjusted q-values.

**S3 Table. Differential abundance analysis results.** This table lists the LinDA q-values.

## Declaration of AI and AI-assisted technologies in the writing process

During the preparation of this work, the authors employed Grammarly (version 1.2.149.1641) to enhance readability and ensure adherence to grammatical and spelling conventions. Subsequently, they meticulously reviewed and made requisite modifications to the content.

## Notes

### Competing Interest Statement

The authors have declared no competing interest.

